# The *macrophage-expressed gene* (*mpeg*) *1* identifies a subpopulation of B cells in the adult zebrafish

**DOI:** 10.1101/836098

**Authors:** Giuliano Ferrero, Etienne Gomez, Sowmya Iyer, Mireia Rovira, Magali Miserocchi, David M. Langenau, Julien Y. Bertrand, Valérie Wittamer

**Affiliations:** Institut de Recherche Interdisciplinaire en Biologie Humaine et Moléculaire (IRIBHM); ULB Institute of Neuroscience (UNI), Université Libre de Bruxelles (ULB), Brussels, Belgium; Department of Pathology and Immunology, University of Geneva, School of Medicine, Geneva, Switzerland; Department of Pathology and Center for Cancer Research, Massachusetts General Hospital Research Institute, Boston, MA, USA; WELBIO, Belgium

**Keywords:** Macrophages, B lymphocytes, Perforin-2, *mpeg1*

## Abstract

The mononuclear phagocytic system (MPS) consists of many cells, in particular macrophages, scattered throughout the body. However, there is increasing evidence for the heterogeneity of tissue-resident macrophages, leading to a pressing need for new tools to discriminate MPS subsets from other hematopoietic lineages. *Mpeg1.1* is an evolutionary conserved gene encoding *perforin-2*, a pore-forming protein associated with host defense against pathogens. Zebrafish *mpeg1.1:GFP* and *mpeg1.1:mCherry* reporters were originally established to specifically label macrophages. Since, more than 100 peer-reviewed publications have made use of *mpeg1.1*-driven transgenics for *in vivo* studies, providing new insights into key aspects of macrophage ontogeny, activation and function. However, while the macrophage-specific expression pattern of the *mpeg1.1* promoter has been firmly established in the zebrafish embryo, it is currently not known whether this specificity is maintained through adulthood. Here we report direct evidence that beside macrophages, a subpopulation of B-lymphocytes is marked by *mpeg1.1* reporters in most adult zebrafish organs. These *mpeg1.1^+^* lymphoid cells endogenously express *mpeg1.1* and can be separated from *mpeg1.1*^+^ macrophages by virtue of their light-scatter characteristics using FACS. Remarkably, our analyses also revealed that B-lymphocytes, rather than mononuclear phagocytes, constitute the main *mpeg1.1*-positive population in *irf8^null^* myeloid-defective mutants, which were previously reported to recover tissue-resident macrophages in adulthood. One notable exception are skin macrophages, whose development and maintenance appear to be independent from *irf8*, similar to mammals. Collectively, our findings demonstrate that *irf8* functions in myelopoiesis are evolutionary conserved and highlight the need for alternative macrophage-specific markers to study the MPS in adult zebrafish.

**SUMMARY SENTENCE:** Mpeg1 is not a restricted macrophage marker, but also labels B cells in the adult zebrafish. Therefore, previously identified *irf8*-independent macrophages likely consist of B lymphocytes.

**Graphical Abstract:** 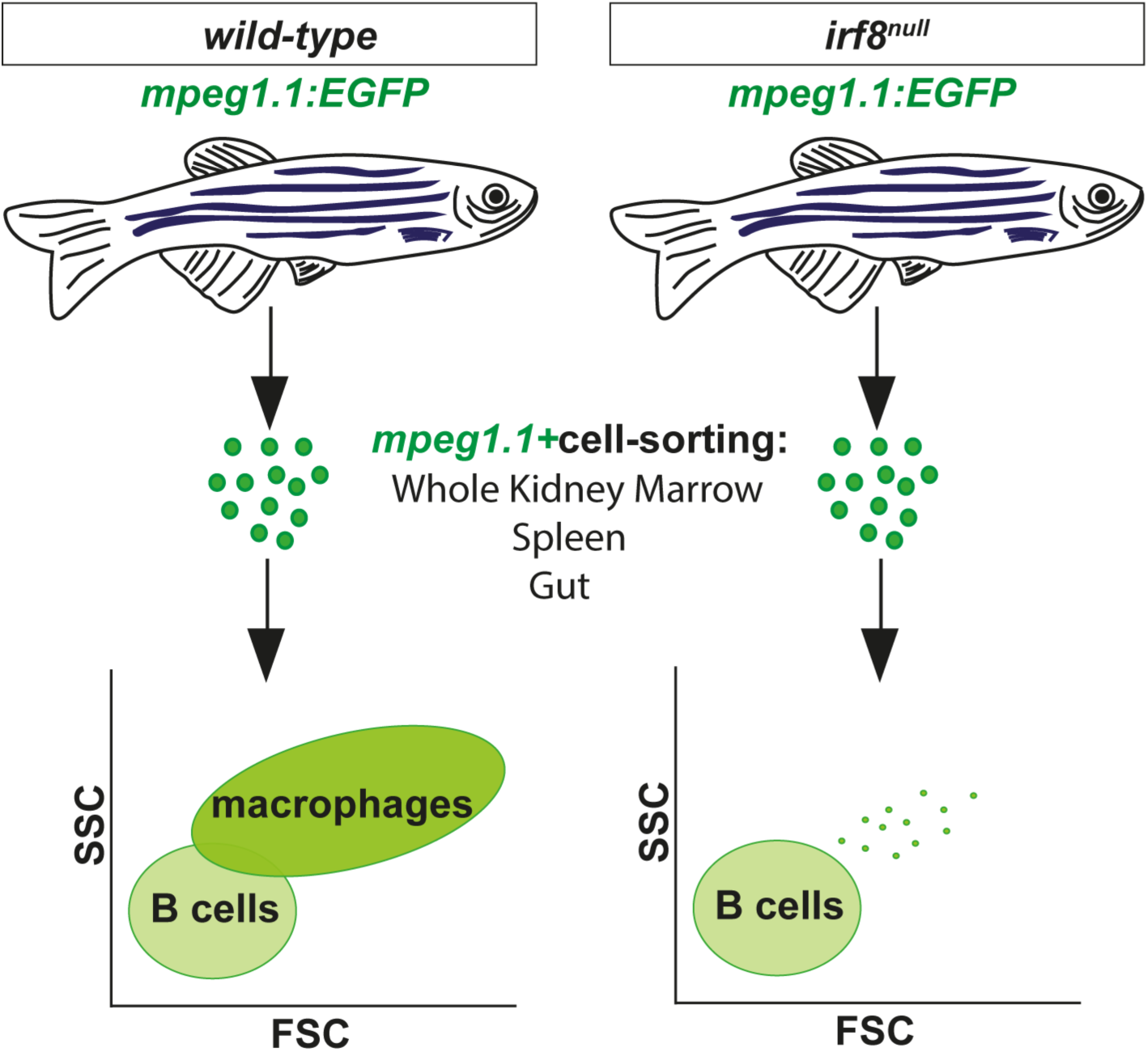

## INTRODUCTION

Over the last decades, the zebrafish has become a powerful model to study biological processes conserved with mammals^1, 2^. In the fields of hematopoiesis and immunology, zebrafish has provided a new level of precision in understanding key aspects of developmental immunity and disease^3, 4^, some of which are currently being tested in the clinic^5^. Consequently, many efforts have been engaged to thoroughly characterize the zebrafish immune system. Using proximal promoter elements of lineage-specific genes, we and others have established fluorescent transgenic lines that specifically mark the major blood cell lineages in the zebrafish^6, 7^. Combined with light scatter separation, these lines have allowed the prospective isolation of zebrafish immune cells by flow cytometry, and their subsequent morphological, phenotypical and functional characterization. Collectively, these studies have highlighted the high degree of conservation between zebrafish and mammalian immune cell types.

Macrophages, which represent an important branch of the innate immune system^8^ and also exhibit key developmental functions^9^, are, together with neutrophils, one of the most studied leukocyte populations in the zebrafish model. This is mainly due to the availability of several reporter lines, among which the *mpeg1.1* promoter-driven fluorescent reporters are the most frequently used^10^. The *mpeg1.1* gene encodes a pore-forming protein named *Perforin-2*, which is highly related to *Perforin-1* and the *C9* sub-unit of the complement^11, 12^. However, while the two latter proteins are only found in vertebrates and act by killing extracellular targets, *Mpeg1/Perforin-2* is already present in early multicellular organisms like sponges, and targets intracellular pathogens^13^. *Mpeg1.1:GFP* and *mpeg1.1:mCherry* reporter fish have been instrumental in characterizing the behavior of macrophages through live imaging in transgenic embryos, and the *mpeg1.1:Gal4* line for analyzing macrophage-targeted gene function. Altogether, these studies have tremendously contributed to increasing our knowledge on the roles of macrophages in multiple processes of developmental physiology^14–16^, as well as in pathological mechanisms involved in human disease, such as inflammation, infection^17, 18^ and cancer^19–21^. However, while most of the field has initially focused on early macrophages taking advantage of the optical transparency of the zebrafish embryo and larvae, a growing number of investigators are now using these lines to address multiple aspects of macrophage biology in adults. This raises important questions, as the specificity of the *mpeg1.1* driver in the adult hematopoietic system still remains to be determined^22^. Indeed, although *mpeg1.1* was originally described as a macrophage-specific gene in mammals^11^, recent evidence demonstrates that its expression is not restricted to mononuclear phagocytes^13^.

In this study, we initially aimed at characterizing different subsets of macrophages in the adult zebrafish, by combining *mpeg1.1* transgenics with available lines marking the blood compartment. These extended analyses revealed a previously unappreciated cellular heterogeneity of the *mpeg1.1*^+^ population, as we identified a subset of *mpeg1.1*-positive cells that are distinct from macrophages. Molecular characterization showed that these cells consisted of B lymphocytes. A detailed examination of their tissue distribution further demonstrated that *mpeg1.1*-expressing B-cells were present in all major adult lymphoid organs in zebrafish. In addition, *mpeg1.1* expression was also found outside the hematopoietic system, in a subset of epithelial cells located in the skin, as recently described^23^. Finally, we show that adult zebrafish deficient for *irf8* recover *mpeg1.1*^+^ cells, in agreement with previous observations^24^. However, based on our findings, we propose that in most tissues recovering *mpeg1.1^+^* cells consist of B lymphocytes and do not represent *irf8*-independent macrophages. One exception is the epidermis, where the presence of *mpeg1.1^+^* macrophages is mildly affected by the absence of *irf8*.

## MATERIAL AND METHODS

### Zebrafish husbandry

Zebrafish were maintained under standard conditions, in accordance with institutional (ULB) and national ethical welfare guidelines and regulations. All experimental procedures were approved by the ULB ethical committee for animal welfare (CEBEA). The following lines were used: *Tg*(*mhc2dab:GFP-LT)^sd67^*; *Tg(mpeg1.1:eGFP)^gl22^*^[10]^; *Tg(mpeg1.1:mCherry)^gl23^*^[10]^; *Tg(kdrl:Cre)^s898^*^[25]^; *Tg*(*actb2:loxP-STOP-loxP-DsR^edexpress^*)*^sd5^*^[25]^; *Tg(Cau.Ighv-ighm:EGFP)^sd19^*^[26]^is referred to as *ighm:GFP*. Homozygous *irf8^std95/std95^* mutants were derived from heterozygous incrosses of *irf8^std95/+^* fish^24^. Special care was taken to control reporter gene dosage through experiments (with all control and mutant animals used in this study known to carry similar hemizygous or homozygous doses of the GFP transgenes). Unless specified, the term “adult” fish refers to animals aged between 3-6-months.

### Flow cytometry and cell sorting

Single-cell suspensions of dissected adult zebrafish organ were prepared as previously described^7^. Flow cytometry and cell sorting were performed with a FACS ARIA II (Becton Dickinson). Analyses were performed using the FlowJo software (Treestar). Cytospins and May-Grunwald/Giemsa stains were performed as previously described^27^.

### Quantitative PCR

RNA extraction from sorted cells and cDNA synthesis were performed as described^28^. Biological triplicates were compared for each subset. Relative amount of each transcript was quantified via the *ΔCt method,* using *elongation-Factor-1-alpha (ef1α)* expression for normalization, or via the *ΔΔCt* method, using *ef1α* and WKM for normalization. Primers are listed in Table 1.

### Immunostaining and confocal microscopy analyses

Guts were dissected from adult fish, fixed in 4% PFA overnight (O/N) and incubated in 30% sucrose:PBS O/N before snap-freezing in OCT. Scales were manually detached from euthanized fish and pre-treated with 100mM DTT before O/N fixation in 4% PFA. Immunostaining on gut cryosections or floating scales was performed as described^28^, using the following primary and secondary antibodies: chicken anti-GFP polyclonal antibody (1:500; Abcam), rabbit anti-DsRed polyclonal antibody (1:500; Clontech), Alexa Fluor 488-conjugated anti-chicken IgG antibody (1:500; Invitrogen), Alexa Fluor 594-conjugated anti-rabbit IgG (1:500; Abcam). Images were taken with a Zeiss LSM 780 inverted microscope, using a Plan Apochromat 20× objective. Image post-processing (contrast and gamma adjust) were performed with the Zeiss Zen Software.

### IPEX collection and phagocytosis assay

Intraperitoneal exudate (IPEX) of adult zebrafish was collected as previously described^29^. For phagocytosis assay, 2,5 µl of FluoSpheres^TM^ 1µm, blue fluorescent (Thermofisher) were injected in the peritoneal cavity, resulting in a cell:beads ratio ranging between 1:50 and 1:100. IPEX was collected at different time points, resuspended in 1% inactivated fetal bovine serum and analyzed by flow-cytometry with a LSRFortessa^TM^ machine (BD Biosciences).

### Single cell transcriptome analysis

Whole-kidney marrows were isolated from three separate adult wild-type fish (3-9-mo-old) and subjected to inDrop single cell RNA sequencing^30^. These data are publicly-available at the NCBI GEO database under accession ID GSE100913 and can be accessed directly through the website https://molpath.shinyapps.io/zebrafishblood/

### Statistical analyses

For Figure 3D, t-test (with multiple-testing correction) was used to derive p-values. All plots and statistical analysis were performed using R^31^.

## RESULTS

### The *mpeg1* transgene marks distinct populations of leukocytes in the zebrafish WKM

In *mpeg1.1:GFP* or *mpeg1.1:mCherry* transgenic adult zebrafish, parenchymal microglia can be isolated from other CNS-associated macrophages by flow cytometry based on fluorophore expression levels^28^. We therefore sought to investigate whether *mpeg1.1* reporters could similarly discriminate distinct macrophage subsets in other tissues. To facilitate our study, we used *mpeg1.1:mCherry*; *mhc2dab:GFP_LT_* animals, where GFP marks antigen presenting cells within the adult hematopoietic tissue^7^. This strategy was expected to specifically co-label mononuclear phagocytes, allowing tracking of these cells according to multiple parameters. Accordingly, we detected double positive cells by flow cytometry in the WKM (Fig. 1). This population could be fractionated into two subsets, based on levels of GFP and mCherry: the double positive^lo^ population (*mpeg1.1^lo^ mhc2dab^+^*; Dp^lo^) and the double positive^hi^ population (*mpeg1.1^hi^ mhc2dab^+/hi^*; Dp^hi^), accounting for approximately 70% and 30% of total double positive cells, respectively. Since mature mononuclear phagocytes were previously identified as a distinct forward scatter (FSC)^hi^ side scatter (SSC)^int^ population, overlapping the conventional myeloid and precursor fractions^7^, we next examined the scatter profile of these two populations. Surprisingly, we found that only the Dp^hi^ cells localized in the myeloid/progenitor scatter fraction, whereas nearly all Dp^lo^ cells belonged to the FSC^lo^SSC^lo^ lymphoid fraction. Because this latter fraction primarily contains lymphocytes^7, 32^, these observations suggested that fluorophore expression in the *mpeg1.1:mCherry* line was not restricted to macrophages only.

**Figure 1.**
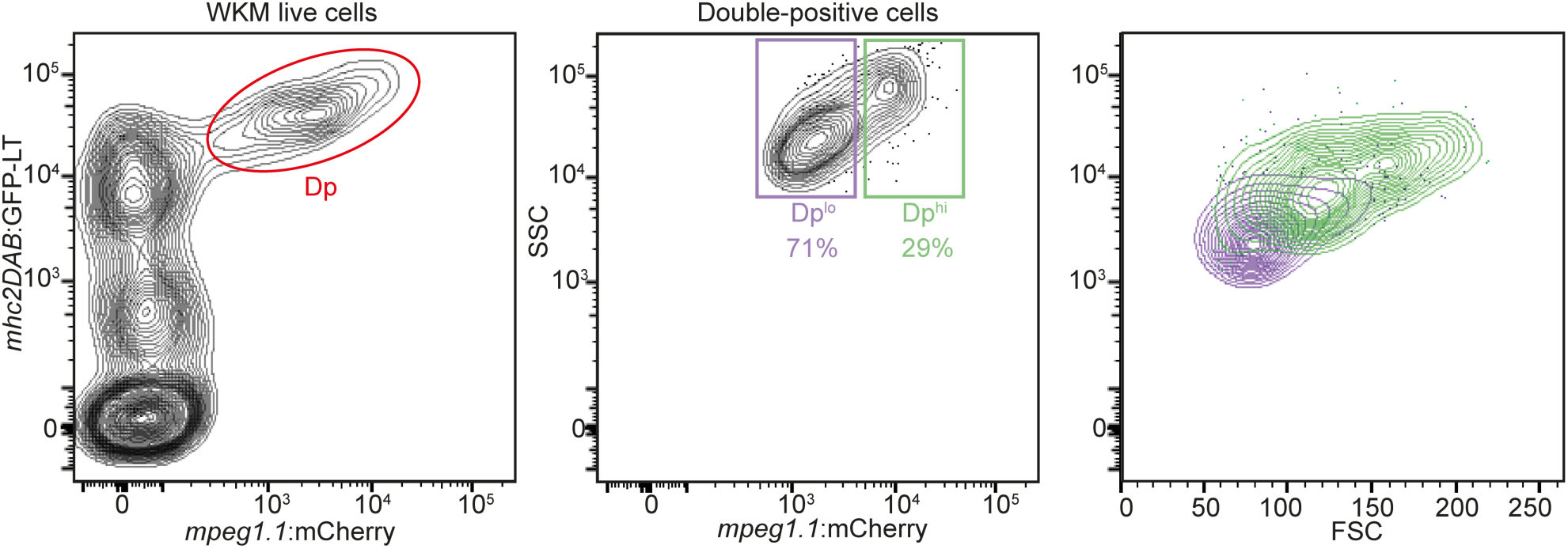
Two populations of *mpeg1.1:mcherry*-positive cells in the adult WKM. The WKM of *Tg*(*mpeg1.1:mcherry*; *mhc2dab:GFP_LT_)* adult reporter fish was dissociated and analyzed by flow-cytometry. The whole *mpeg1.1*^+^; *mhc2dab:GFP*^+^ double-positive (Dp) population is circled in red (left panel) and is further subdivided into a *mpeg1.1^lo^;mhc2dab^lo^* (Dp^lo^) and a *mpeg1.1^hi^;mhc2dab^hi^* (Dp^hi^) fraction based on fluorescence intensity (middle panel). Right panel: separation of the Dp^hi^ and Dp^lo^ populations by light-scatter characteristics. SSC= side scatter; FSC= forward scatter. Percentages of each population refer to a single individual and are relative to the total Dp population (mean ± SD of 3 fish: see text).

To evaluate whether expression within the WKM lymphoid gate was a common feature of all *mpeg1.1* transgenics, we next turned to *mpeg1.1:GFP* adult fish. Flow cytometry analysis revealed that, similar to their *mCherry* counterparts, *mpeg1.1:GFP*-labeled cells represented approximately 13% of the WKM (data not shown). This prompted us to examine in more details GFP expression among the different hematopoietic subsets (Fig. 2Ai). Within the combined myeloid and progenitor fractions, *mpeg1.1:GFP*^+^ cells accounted for 4.0 ± 0.3 % of total cells (Fig. 2Aii, red gate), in accordance with our previous estimations of mononuclear phagocytes abundance in this tissue^7^. As expected, GFP was not detected within the eosinophil fraction (FSC^hi^SSC^hi^; data not shown). However, and validating our initial observations in *mpeg1:mCherry* animals, the analysis of the lymphoid fraction revealed the presence of a prominent *mpeg1:GFP^lo^* population, accounting for 53.2 ± 8.3 % of cells within the lymphoid gate (n=3) (Fig. 2Aiii, green gate). We conclude that the *mpeg1* reporters show a previously unappreciated expression pattern within the lymphoid lineage in the WKM.

### The lymphoid *mpeg1*-positive cells are B-cells

Since we initially observed that lymphoid *mpeg1:GFP^+^* cells co-expressed the antigen-presenting cell *mhc2dab* transgene, we hypothesized they may be B lymphocytes^7^. We sorted GFP^+^ and GFP^-^ subsets from the WKM lymphoid fraction of *mpeg1.1:GFP* fish and scored them for hematopoietic markers by qPCR, comparing them to GFP subsets isolated from the combined myeloid and precursor fractions (hereafter referred to as the PM fraction) (Fig. 2Ai, 2B). Consistent with their expected monocyte/macrophage identity, GFP^+^ PM cells showed strong expression of *mpeg1.1* and *csf1ra*, but not *mpx*, a neutrophil marker (Fig. 2Bi). Within the lymphoid gate, GFP^+^ cells also expressed *mpeg1.1*, although at a lower level. Expression of *mpeg1.1* mRNA was undetectable in the lymphoid GFP^-^ population, thus indicating that fluorophore expression in the *mpeg1* reporter line reflects endogenous gene expression. Interestingly, lymphoid GFP^+^ cells did not express *csf1ra* (Figure 2Bi), supporting the hypothesis that this population represents a leukocyte subset distinct from mononuclear phagocytes. Rather, examination of canonical B-cell-associated genes revealed strong expression of *pax5, ighz* and *ighm* (Fig. 2Bii). The expression of B-cell genes was not restricted to *mpeg1:GFP^+^* cells within the lymphoid fraction, as similar levels of transcripts were also found in the GFP^-^ subset. Additionally, the lymphoid GFP^+^ cells also lacked expression of T-cell markers such as *lck*, *cd28* and *cd4.1*, which were found to be highly enriched in the GFP^-^ subset (Fig. 2Biii). Altogether, these results indicate that *mpeg1:GFP* marks a subpopulation of B lymphocytes that contains a mixture of *IgM*- and *IgZ*-expressing cells. We concluded that, in addition to mononuclear phagocytes, *mpeg1.1* also marks cells from the B-cell lineage. These findings were further supported by imaging using May-Grünwald-Giemsa staining, showing that the majority of lymphoid *mpeg1:GFP^+^* cells exhibit the expected morphology of lymphocytes, while PM GFP^+^ cells harbored features of macrophages (Fig. 2C), as previously described^7, 32^.

**Figure 2.**
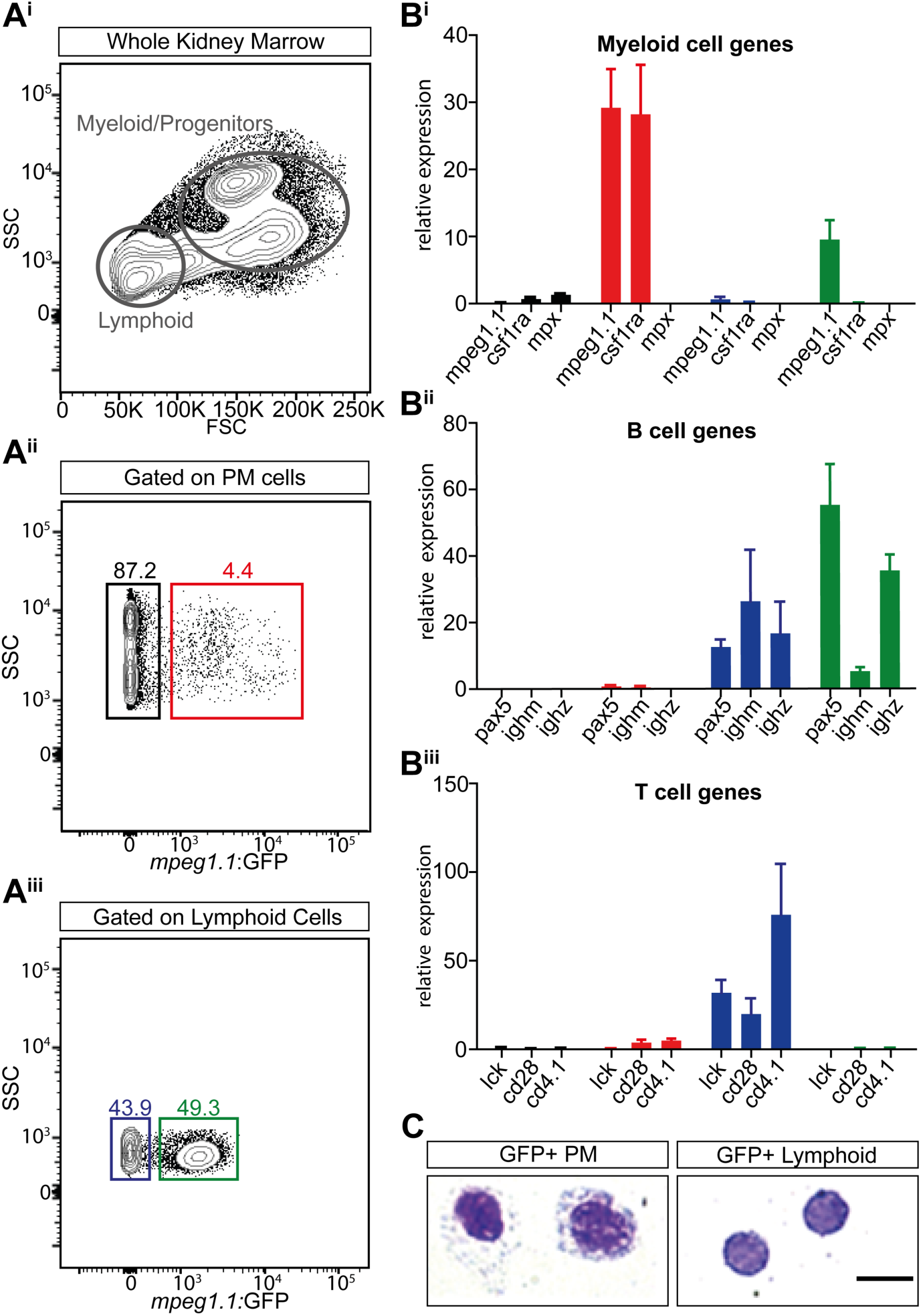
*Mpeg1.1* expression identifies a subset of B-cells in the adult WKM. (A) Gating strategy to isolate lymphoid and myeloid-progenitors (MP) cells from the WKM using light-scattering characteristics (A^i^). Expression of *mpeg1.1:GFP* in the MP (A^ii^) and lymphoid (A^iii^) fractions. Throughout the figure, the GFP^-^ MP fraction is denoted by a black gate and bars, *GFP^+^* MP by a red gate and bars, *GFP^-^* lymphoid by a blue gate and bars and *GFP^+^* lymphoid by a green gate and bars. Percentages represent a single individual and are relative to the total live cells (mean ± SD of 3 fish: see text) (B) Q-PCR expression for genes specific to the myeloid (B^i^), B-(B^ii^) and T-(B^iii^) cell lineages in sorted *mpeg1.1:GFP^+^* and *mpeg1.1:GFP^-^* cells. Units on the y-axis represent changes (fold) above WKM, which is set at 1.0. Error bars indicate SEM (n=3). (C) Morphology of *mpeg1.1:GFP^+^* cells isolated from the lymphoid and MP fractions in WKM. Cells were cytospun and stained with May-Grunwald-Giemsa. Myeloid *GFP^+^* cells show the characteristics of macrophages, with kidney-shaped nuclei and vacuoles, while lymphoid *GFP^+^* cells revealed a typical lymphocytic morphology, with a nonlobed nucleus and a high nuclear:cytoplasmic ratio. Images were taken with a Zeiss AxioImager Z1 micorscope, using a 100x oil-immersion objective. Scale bar: 20 µm.

### Single cell analyses of *mpeg1*-expressing WKM cells

Our results revealed the existence of *mpeg1*.*1^+^* B lymphocytes in the adult WKM. However, B-cell transcripts were also found in the lymphoid *mpeg1.1*^-^ cell population, suggesting that not all B lymphocytes express *mpeg1.1.* To further explore the identity of lymphoid *mpeg1.1^+^* cells, we turned to single cell analysis and examined transcript expression among individual hematopoietic cells using our previously reported single cell transcriptional profiling of zebrafish WKM^30^. By querying this dataset, we first confirmed the existence of a *mpeg1.1^+^* cluster associated with the B-cell lineage (Fig. 3A,B). Extending our initial findings, we further found that these cells co-expressed a large array of B-specific genes including *cd79a*, *cd37*, *ighm*, *pax5,* as well as several transcripts encoding different light chain isotypes (Fig. 3C). As expected, B-cell gene expression was negligible in cell clusters positive for the macrophage transcripts *marco* or *mfap4* (Fig. S1). From these analyses, we estimated that *mpeg1.1^+^* cells accounted for approximately 28.5% of total B lymphocytes (N=47 out of 165 B-cells found in the WKM), in line with our cytometry analysis. Next, we performed a comparative expression analysis between WKM-derived *mpeg1.1^+^* B-cells and macrophages. Violin plots show the distribution of cells expressing each transcript (Fig. 3D). In agreement with the levels of fluorophores observed by flow cytometry, *mpeg1.1* was expressed slightly higher in macrophages than in B-cells. As expected, *marco* and *mfap4* belonged to genes that were uniquely expressed in *mpeg1.1^+^* macrophages, whereas *cd79a*, *ighm* and *ighd* were expressed significantly higher in *mpeg1.1^+^* B-cells. *Ighz* and *pax5* were also more expressed in B-cells than in macrophages, but the very low cell number did not allow to reach significance. Collectively, these unbiased analyses of single cell gene expression showed that lymphoid *mpeg1.1*^+^ cells display a robust B lymphocyte signature.

**Figure 3.**
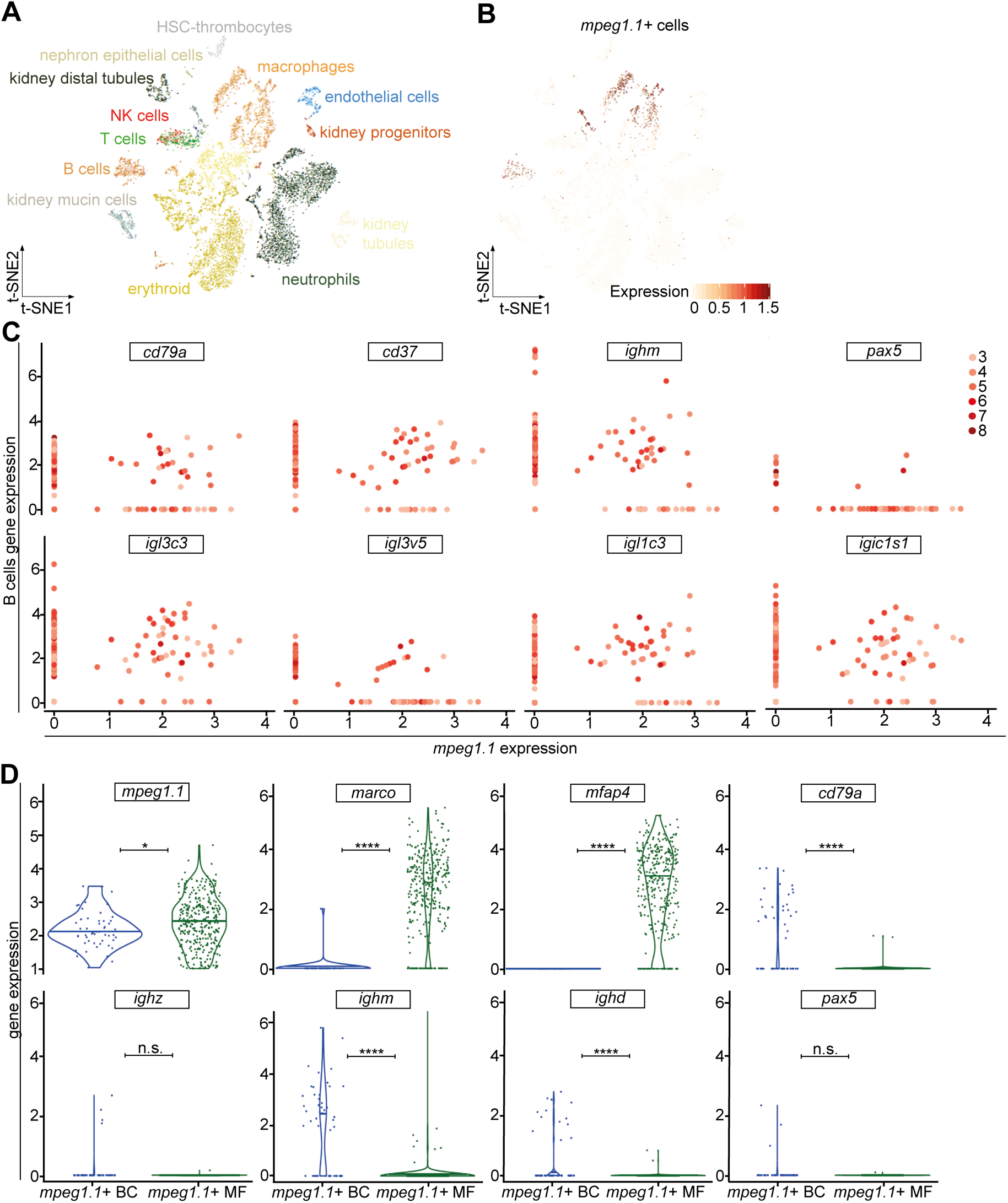
Characterization of *mpeg1.1* expression in adult hematopoiesis by single cell RNAseq analysis. (A) 2D projection of the tSNE analysis showing the hematopoietic and non-hematopoietic cell types of the adult WKM, identified by single-cell InDrops RNA sequencing. (B) Analysis of *mpeg1.1* expression (red) across the clusters in the tSNE plot. Intensity of the color is proportional to the expression level. (C) Log of normalized and scaled expression of B-cell genes (y-axis) and *mpeg1.1* (x-axis) in single-cells from the WKM. Each scatter-plot represents the expression of a B-cell gene, indicated on top. Dots correspond to individual cells and the color-code on the right describes the overall number of B-cell markers expressed by each cell. (D) Violin plot analysis comparing gene expression (y-axis) between *mpeg1.1*^+^ macrophages and B-cells.

### Tissue distribution of *mpeg1.1*^+^ B-cells

Since we demonstrated that *mpeg1.1^+^* lymphocytes and myeloid cells could be discriminated by their expression of *ighm*, we next crossed *mpeg1.1:mCherry* fish to *ighm:GFP* transgenic animals, which specifically mark *IgM*-expressing B lymphocytes^26^. As expected, flow cytometry analysis of adult double transgenic fish identified a prominent double positive cell subset in the WKM (Fig. 4Ai). This population, defined as *mpeg1.1^lo^ighm^+^*, accounted for 11.5 ± 7.4 % of total WKM blood cell numbers (n=6). Two other subsets with differential levels of *mCherry* and *GFP* expression (*mpeg1.1^hi^ighm^-^* and *mpeg1.1^-^ighm^hi^*) were also present, although less abundant (4.2 ± 0.7 % and 4.2 ± 1.7 % of total WKM cell numbers, respectively; n=6). Based on our previous gene expression analyses, the *mpeg1.1^+^ighm^-^* population likely contains a mixture of mononuclear phagocytes and *mpeg1.1^+^ IgZ*-expressing B lymphocytes, which are not labeled in the *ighm:GFP* line^26^. The fractionation of *ighm:GFP^+^* cells into two subpopulations according to *mpeg1.1* expression was also consistent with our initial identification of *ighm* transcripts in both *mpeg1.1:GFP^+^* and *mpeg1.1:GFP^-^* lymphoid subsets, and reflects the heterogeneity of the *ighm:GFP^+^* population. Importantly, *mpeg1.1^+^ ighm^+^* B-cells *(mpeg1^lo^ighm^+^*) appeared to constitute the prominent cell subset among *IgM-*expressing cells, as approximately 70% of GFP^+^ cells co-expressed *mpeg1.1:mCherry*.

We next sought to determine whether the two subsets of B-cells existed in peripheral tissues. Using the same *mpeg1.1:mCherry*;*ighm:GFP* double transgenic fish, we analyzed the spleen, gut and skin, known to host *IgM*-expressing B lymphocytes^26, 33^. While the proportion of *IgM*^+^ cells greatly varied among tissues, the majority of *IgM^+^* cells showed co-expression with *mpeg1.1* in all samples examined (Fig. 4A). Confocal imaging and immuno-histochemistry analyses confirmed the presence of double positive cells scattered in the skin or located along the basal lamina propria of the gut in double transgenic animals (Fig. 4B). Interestingly, the skin double positive cells displayed a dendritic morphology that was barely distinguishable from single *mpeg1.1^+^ighm^-^* cells, probably due to the fact that all these cells are located between epithelial cells in the epidermis. Collectively, these results indicate that *mpeg1.1* expression is persistent in mature B lymphocytes and labels nearly all *IgM^+^* B-cells in the periphery.

**Figure 4.**
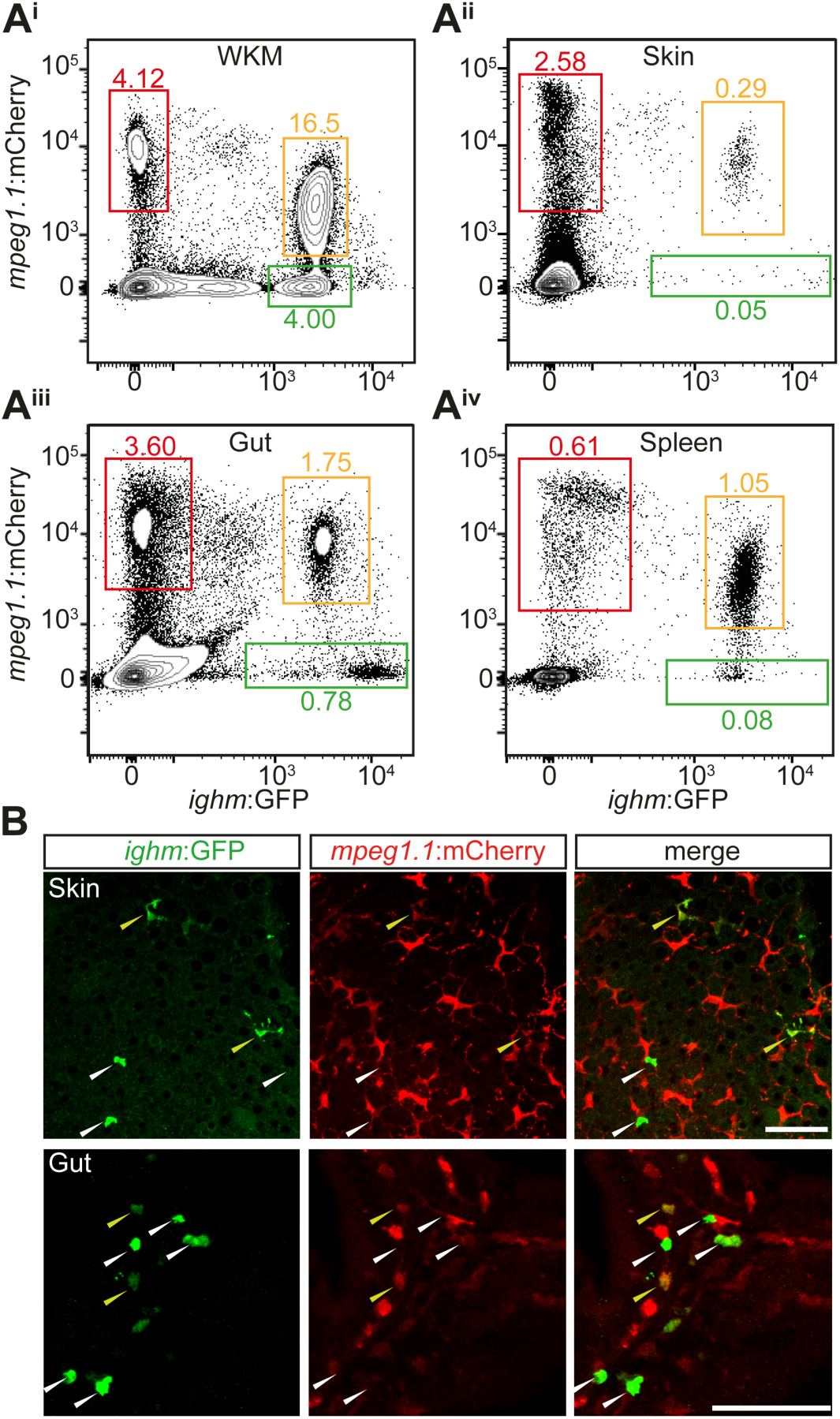
Distribution of *mpeg1.1*-positive B-cells in adult tissues. (A) Distribution of fluorescence in cell suspensions prepared from WKM (A^i^), skin (A^ii^), gut (A^iii^) and spleen (A^iv^) of an adult *Tg*(*ighm:GFP*; *mpeg1.1:mCherry*) fish. In each organ, three main populations can be discriminated: *mpeg1.1^hi^; ighm^-^* cells (red gate); *mpeg1.1^-^; ighm^hi^* B cells (green gate); and *mpeg^lo^*; *ighm^+^* cells (orange gate). Percentages represent the single individual (mean ± SD of 5 fish: see text). (B) Immunofluorescence on skin scales (upper panels) and gut sections (lower panels) from an *Tg*(*ighm:GFP*; *mpeg1.1:mCherry*) reporter fish. The GFP (left), mCherry (middle) and merged (right) images are shown. White arrowheads indicate *ighm^+^; mpeg1.1^-^* cells; yellow arrowheads indicate *ighm^+^; mpeg^+^* cells. Images were taken with a Zeiss LSM 780 inverted confocal microscope, using a 20x plan apochromat objective. Scale bar: 50µm.

### Mpeg1 expression does not discriminate a subset of phagocytic B cells in zebrafish

Previous studies have established the central role of *mpeg1* in killing intracellular bacteria^13^, as well as the phagocytic capabilities of teleost B cells^34^. Given the clear distribution of *ighm^+^* B cells into *mpeg1.1^+^* and *mpeg1.1^-^* populations, we thus next wondered whether expression of the microbicidal *mpeg1.1* gene could discriminate a subset of phagocytic B cells. We injected 1µm blue-fluorescent FluoSpheres in the peritoneal cavity of *mpeg1.1:mCherry*; *ighm:GFP* adult fish, where *mpeg1.1*+ B cells are present in steady-state conditions (Figure 5A). We then collected the intraperitoneal exudate (IPEX) at different time points and quantified the percentage of phagocytic cells within each fraction by flow-cytometry. As expected, both the *mpeg1.1^+^ighm^-^* and *mpeg1.1^-^ighm^-^* cell populations, which contain professional phagocytes (macrophages and polynuclear cells, respectively) actively engulfed blue beads between 2 and 8 hours post-injection (hpi) (Figure 5B). By contrast, the phagocytic capacity of *ighm^+^* B cells was negligible, as we did not observe any particle internalization for either *mpeg1.1^+^* or *mpeg1.1^-^ ighm^+^* B cells (Figure 5B). Importantly, all phagocytic *ighm^-^* cells clustered within the myeloid gate (data not shown), therefore ruling out the presence of *ighm^-^* B cells in the phagocytic population. Together, these results indicate that zebrafish *ighm^+^* B cells lack phagocytic activity, in line with our previous observations in this model^26^.

**Figure 5.**
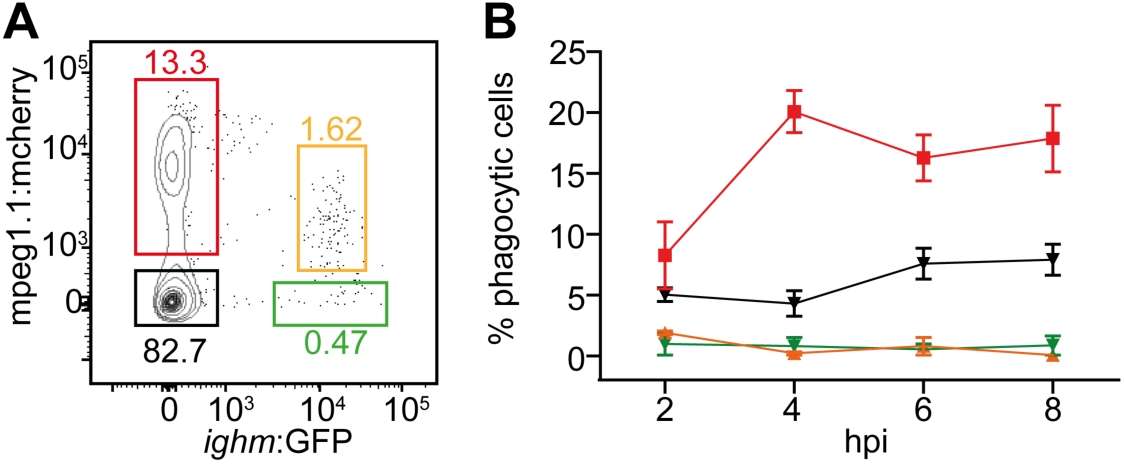
*Mpeg1.1* expression does not discriminate a subset of phagocytic B cells in zebrafish. (A) Distribution of fluorescence in cell suspension from the IPEX of adult *Tg*(*ighm:GFP*; *mpeg1.1:mCherry*) fish. Percentages of cells in each gate refer to the total live cell fraction and represent an average of 3 individuals. (B) Percentage of phagocytic cells within each of the fractions described in (A), at different hours post-injection (hpi) of blue fluorescent beads. Each time point represents the average of 3 individuals.

### Adult tissue macrophages, except in the skin, develop in an *irf8-*dependent manner

The unexpected observation that B lymphocytes represent a significant proportion of the total *mpeg1.1:GFP^+^* population in adults prompted us to examine in more details the phenotype of macrophage-deficient mutants previously characterized using *mpeg1.1* reporters. Notably, we focused on mutants for the macrophage-specific transcription factor *irf8*, as they are devoid of mature macrophages during embryonic life, but seem to recover later in life, based on the presence of *mpeg1.1:GFP^+^* cells in juvenile and adult fish^24^. To rigorously assess the lineage identity of *irf8*-independent *mpeg1.1^+^* cells, we analyzed hemato-lymphoid organs isolated from *irf8^null^;mpeg1.1:GFP* adult animals by flow cytometry. In the WKM, *GFP^lo^* B lymphocytes were present in proportion similar to that of their *wild-type* siblings (Fig. 6A, red gate). However, the mutants exhibited a profound deficit in macrophages, as demonstrated by the complete loss of the minor *GFP^hi^* population (Fig. 6A, blue gate). In the periphery, examination of spleen, gut and skin cell suspensions showed no reproducible alterations in the frequency of *mpeg1.1*^+^ cells. However, flow cytometry profiling revealed important differences between controls and *irf8^null^* fish (Fig. 6B,C). When back-gated to the lymphoid or myeloid lineages, we found, as expected, that the GFP^+^ populations in control animals scattered through both the lymphoid and myeloid gates (Fig. 6B). In contrast, the majority of the GFP^+^ cells in the spleen and gut of *irf8^null^* mutants segregated in the lymphoid gate only, thus indicating a deficit in macrophages. However, *irf8^null^ mpeg1.1*^+^ cells in the skin segregated in both the lymphoid and myeloid fractions with frequencies that mirrored the *wild-type* controls, thus suggesting a normal mononuclear phagocyte compartment.

**Figure 6.**
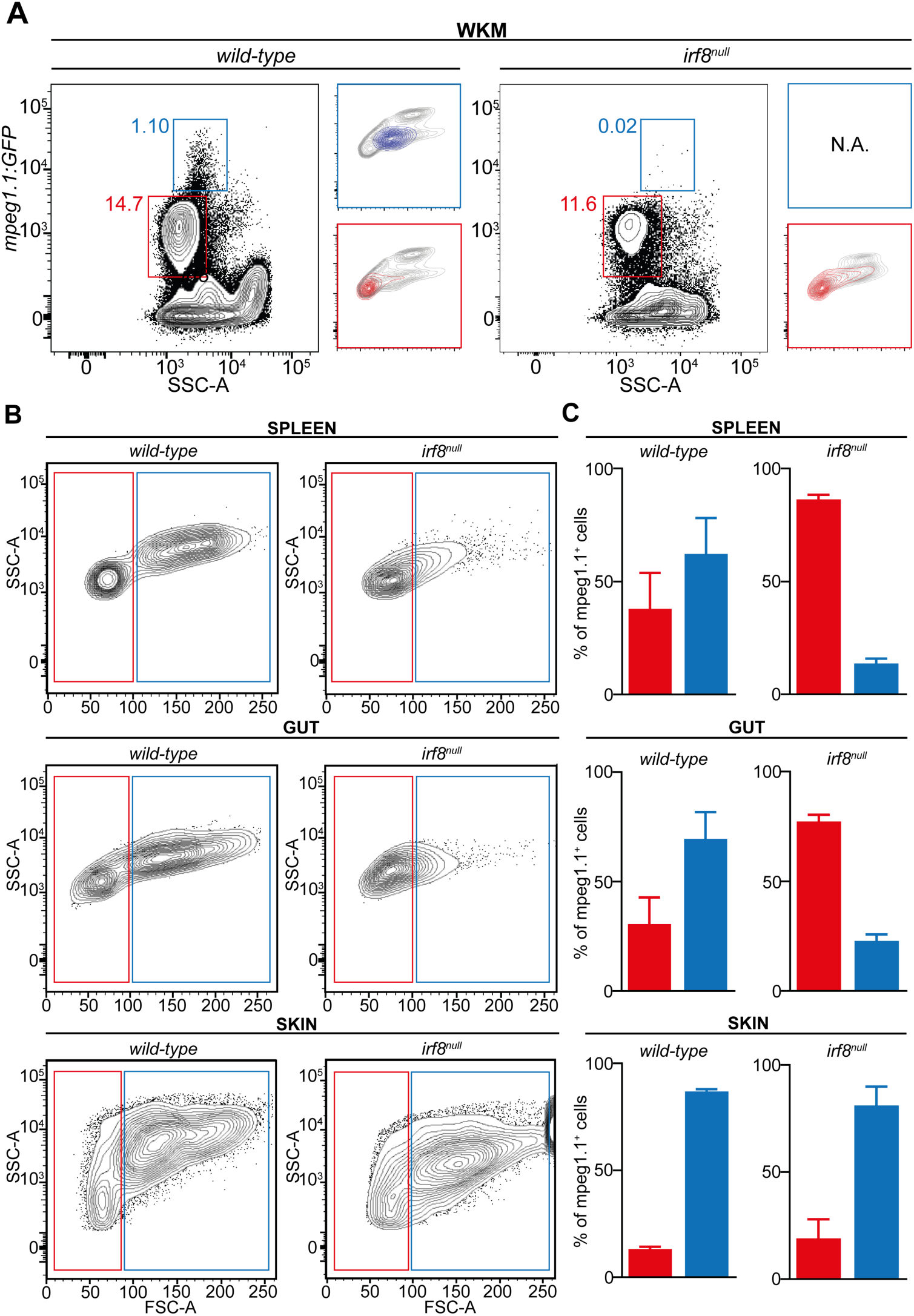
Adult zebrafish *irf8* mutants display myeloid deficiencies. (A, B) Flow-cytometry analysis of organ cell suspensions from adult *wild-type* and *irf8*^null^ fish carrying the *mpeg1.1:GFP* reporter. (A) The whole *mpeg1.1:GFP^+^* cell cluster of the WKM is split into an *mpeg1.1^lo^* (red frame) and a *mpeg1.1^hi^* (blue frame) subsets. Lateral panels show the projection of the *mpeg1.1^lo^ and mpeg1.1^hi^* clusters into the FSC/SSC plot. Percentages relative to the total *mpeg1.1:GFP^+^* cells are indicated aside of each gate and refer to a single individual (mean ± SD of 3 fish: see text). (B) FSC/SSC plots showing the distribution of *mpeg1.1:GFP^+^* cells from the spleen, gut and skin in the lymphoid (red) and MP (blue) gates. (C) Quantification of *mpeg1.1:GFP^+^* lymphoid (red) and MP (blue) cells depicted in (B). Columns represent the percentage relative to the total *mpeg1.1:GFP^+^* population of each organ. Error bars represent SD (n=3).

Because the identification of a distinct *mpeg1.1*^+^ cell population with a non-hematopoietic origin has been recently reported in the zebrafish epidermis, we next sought to evaluate to which extend these cells, referred to as metaphocytes^23^, contributed to the *mpeg1.1*^+^ population found in the skin of *irf8^null^* mutants. To do so, we took advantage of Tg(*kdrl:Cre*;*ßactin2:loxP-STOP-loxP-DsRed*), a fate-mapping model where constitutive Cre expression is targeted to endothelial cells^25^, resulting in permanent labeling of hemogenic endothelium-derived adult hematopoietic cells (Fig. 7A). Due to their distinct ectodermal origin^23^, *mpeg1.1*^+^ metaphocytes were not expected to be labeled in this system. We thus crossed the *irf8* mutant line to Tg(*kdrl:Cre*;*ßactin2:loxP-STOP-loxP-DsRed;mpeg1.1:eGFP*), and analyzed the adult skin for *GFP* and *DsRed* using flow cytometry. In line with our hypothesis, in WT controls, two major *mpeg1.1:GFP^+^* populations could be observed: *GFP^+^DsRed^-^* and *GFP^+^DsRed^+^* (Fig. 7B). To further characterize these fractions, we analyzed them for hematopoietic and epithelial markers (Fig. 7C). Using these criteria, we concluded that the *GFP^+^ DsRed^-^* cells were non-hematopoietic, as demonstrated by the complete lack of expression of the pan-leukocytic *cd45* and other myeloid and lymphoid genes. Rather, *GFP^+^ DsRed^-^* cells expressed high level of *cldnh1* transcripts and likely qualified as the so-called metaphocytes (Fig. 7C). As expected, *mpeg1.1*^+^ mononuclear phagocytes and B-cells belong to the *GFP^+^DsRed^+^* positive population, which was negative for *cldnh1* expression. As shown in Fig. 6B, the relative representation of both populations was not significantly changed in *irf8^null^* fish skin. This indicated that neither the ontogeny or maintenance of metaphocytes rely on *irf8*. In addition, these results also validated our previous observations that the *irf8* deficiency does not affect skin macrophage development in zebrafish. These data were corroborated by fluorescent confocal analyses performed on fish scales (Fig. 7D). Collectively, these findings demonstrated that *irf8* mutant fish have a profound deficit of mononuclear phagocytes in the WKM, spleen and gut but sub-normal numbers of mononuclear phagocytes in the skin. We conclude that, with the exception of the skin, *mpeg1.1^+^* hematopoietic cells in most organs mainly consist of B lymphocytes, which development is largely independent on *irf8*.

**Figure 7.**
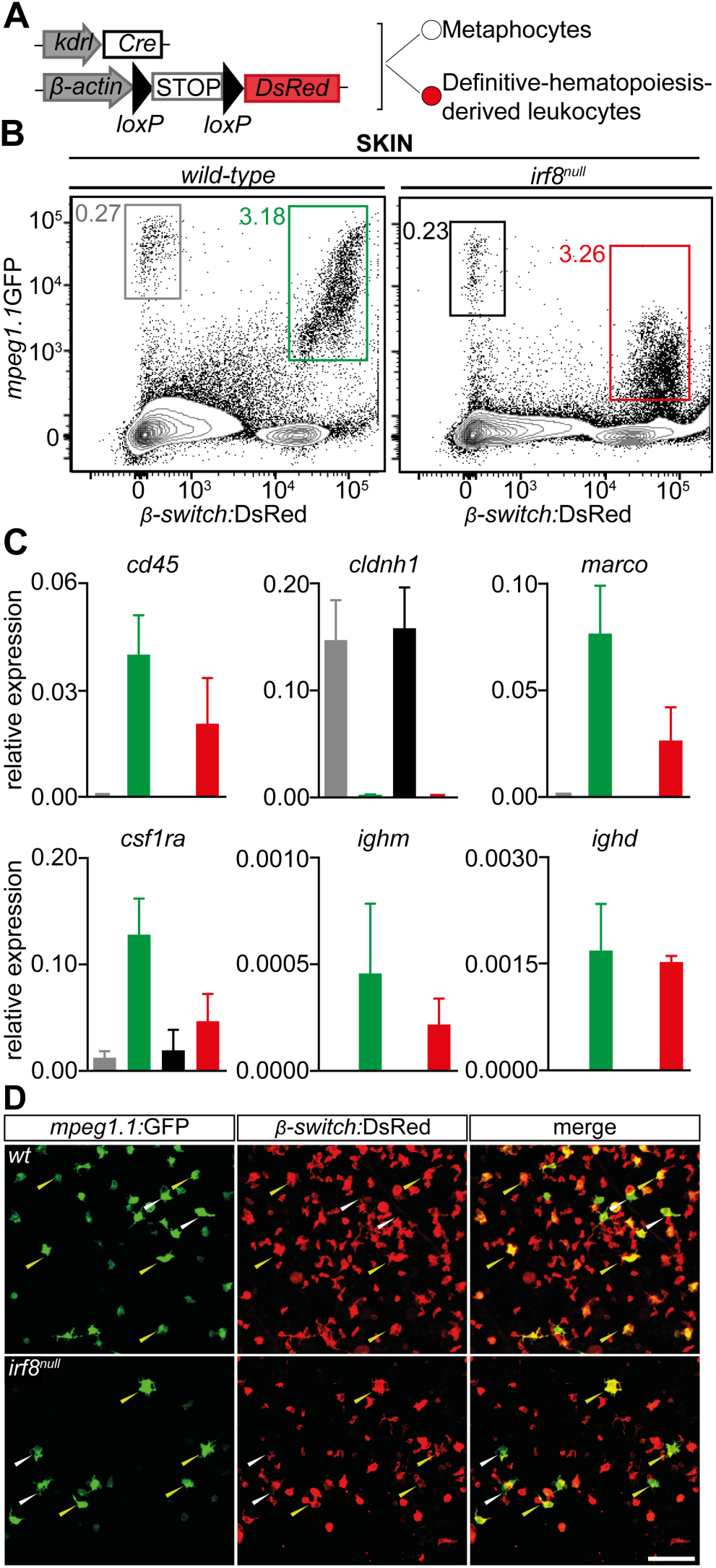
Skin adult macrophages develop independently of *irf8.* (A) Scheme of the transgenic lines used to mark endothelial-derived definitive hematopoiesis (B) Flow cytometry analysis on a skin cell suspension from an adult Tg(*kdrl:Cre*; *β-actin:switch-DsRed*; *mpeg1.1:GFP)* triple transgenic (left panel) discriminates two *mpeg1.1^+^* populations based on *DsRed* fluorescence levels: *mpeg1.1:GFP^+^*; *β-actin:switch-DsRed^-^* (grey gate) and *mpeg1.1:GFP^+^*; *β-actin:switch-DsRed^+^* (green gate) cells. The frequency of each population is unchanged in *irf8*^null^ fish (right panel, grey and red gates). (C) For each genotype, gene expression in *mpeg1.1:GFP^+^*; *β-actin:switch-DsRed^-^* and *mpeg1.1:GFP^+^*; *β-actin:switch-DsRed^+^* cells examined for the presence of hematopoietic (*cd45*), epithelial (*cldnh1*), macrophage (*marco*, *csf1ra*) and B-cell (*ighm*, *ighd*) -specific transcripts. The color-code matches the FACS plots. Units on the y-axis represent transcript expression normalized to *ef1α* expression levels. Error bars represent SEM (n=3). (D) Fluorescence of GFP (left panels) and DsRed (middle panels) in the skin of adult *Tg*(*mpeg1.1:GFP*; *β-actin:switch-DsRed^+^*) *wild-type* (upper panels) and *irf8*^null^ (lower panels) fish. The right panels show a merge of both fluorescent channels (n=3). Yellow arrowheads indicate *mpeg1.1:GFP^+^; β-actin:switch-DsRed^+^* cells; white arrowheads indicate *mpeg1.1:GFP^+^; β-actin:switch-DsRed^-^* cells. Images were taken with a Zeiss LSM 780 inverted confocal microscope, using a 20x plan apochromat objective. Scale bar: 50µm.

## DISCUSSION

The adult zebrafish has recently emerged as a promising model system to perform immunological investigations. However, as a relatively latecomer to the field of immunology, only a limited number of antibodies are available against hematopoietic cells, thus precluding their prospective isolation using techniques traditionally performed in the mouse model. To overcome these limitations, most studies in adult zebrafish take advantage of transgenic lines marking leukocytes. Among these, many have employed *mpeg1.1*-driven transgenes to be a specific reporter for adult macrophages. However, these macrophage-reporter lines were originally created to perform live-imaging studies in transparent embryos and larvae, and their reliability in adults has never been evaluated.

In this work, we disprove the wide assumption that *mpeg1.1* serves as a specific pan-macrophage marker in the adult zebrafish, as we found that B-cells also express *mpeg1.1*. Importantly, we showed that fluorophore expression driven by the 1.2kb-promoter recapitulates the endogenous expression pattern of *mpeg1.1* in B-cells, thus resulting in labeling of *mpeg1.1^+^* B lymphocytes *in vivo,* alongside mononuclear phagocytes. Interestingly, our findings correlate with current knowledge of the *Mpeg1* expression in mammals. Indeed, although *Mpeg1* was originally described in human and mouse as a macrophage-specific gene (hence its name, *<u>m</u>acro<u>p</u>hage-<u>e</u>xpressed <u>g</u>ene <u>1</u>*)^11^, recent evidence indicated that it is more broadly expressed that initially thought. For example, *Mpeg1* was shown to be constitutively expressed in splenic marginal zone (MZ) B cells, where it stands as a specific marker to differentiate them from follicular (FO) B-cells^35^. Similarly, we identified two subsets of B-cells based on *mpeg1.1* expression in zebrafish. However, histological examination of secondary lymphoid tissues did not allow to identify any specific anatomical areas where these two populations would segregate. As teleosts lack well-organized lymphoid structures and do not form germinal centers (a feature that arose in birds across the vertebrate phylum^36^), whether mpeg1.1 can discriminate different zebrafish B cell subsets based on their locations thus remains an open question. In addition, *Mpeg1* expression in mammals was also reported in a large array of innate immune cells, including dendritic cells, neutrophils and NK cells^13^. Based on our analyses in zebrafish, we conclude that *mpeg1.1* is not expressed in neutrophils, nor in other lymphocytes. Interestingly, zebrafish possess three *mpeg1* paralogues^37^ and it remains possible that either *mpeg1.2* or *mpeg1.3*, whose expression pattern is poorly characterized, will mark other hematopoietic populations. Finally, it was shown that in inflammatory conditions, *Mpeg1* can also be induced in non-hematopoietic cells, such as barrier cells (keratinocytes and mucosal epithelia)^13^. While our study did not address whether expression of zebrafish *mpeg1.1* was similarly induced upon inflammation, we did find both endogenous and transgenic expression of *mpeg1.1* outside of the hematopoietic tissue in steady-state conditions, in a subset of cells present in the adult skin. Based on their non-hematopoietic identity and their expression of *cldnh*, these cells likely represent metaphocytes, ectodermal-derived myeloid-like cells with antigen-presenting properties that were recently described in the zebrafish epidermis^23^. However, in this study, the authors reported metaphocytes to represent 30% of the whole *mpeg1.1^+^* population present in the skin, which is not concordant with our findings where only 8% of total *mpeg1.1^+^* cells are non-hematopoietic. This underestimation is likely due to the nature of the tissue sample being analyzed, as our quantification refers to the whole skin (consisting of the three layers: epidermis, dermis and hypodermis) while Lin et al. analyzed the epidermis only^23^.

Our work also provides strong evidence that zebrafish *irf8* is vital for the proper development of macrophages throughout life and identified most *mpeg1.1^+^* hematopoietic cells found in adult *irf8* mutants as B lymphocytes. These results challenge a previous report that concluded, based on the progressive, albeit incomplete, recovery of *mpeg1.1^+^* cells in *irf8* mutant fish, that *irf8* was required for the ontogeny of embryonic macrophages but dispensable for the formation of adult macrophages^24^. In this study, however, the assignment to the macrophage identity exclusively relied on the detection of *mpeg1.1:GFP^+^* cells by fluorescent imaging and was not supported by expression data. Here, by performing a detailed phenotypic characterization of these cells, we unambiguously demonstrated their B-cell identity. Such conclusions are further supported by the fact that the timing of appearance of *irf8*-independent *mpeg1.1^+^* cells (starting at the juvenile stage) coincides with that of the ontogeny of B cells, which occurs at around three weeks of development^26^.

Collectively, our data demonstrated that adult zebrafish *irf8* mutants are devoid of most tissue macrophages, a phenotype that is consistent with the well-established role of *Irf8* as a critical regulator of myelopoiesis in mammals^38^. One exception is the skin, where we found *mpeg1.1*-positive cells harboring a myeloid gene signature. We postulate that these cells may constitute the equivalent of mammalian Langerhans cells, whose development and maintenance appears to be independent of *Irf8* in both humans and mice^39, 40^. However, further characterization of their cytochemical properties, ultrastructure, and additional functions will be required to test whether these cells are the true counterparts of mammalian Langerhans cells. In addition, by excluding a role for *irf8* in the regulatory network that controls the ontogeny and maintenance of metaphocytes, this study also provides valuable insights into the biology of this newly discovered cell population.

Finally, our work sheds new lights on B cell immunity in zebrafish and may open new perspectives in the field of comparative immunology. As expression of perforin-2 in professional phagocytes is tied to microbicidal activity^12, 13, 37, 41^, it was tempting to speculate that *mpeg1* could discriminate functionally distinct B cell populations in the zebrafish. This hypothesis was further supported by evidence from the literature that B cells in teleosts are capable to phagocytose particles and microorganisms and also display microbicidal activity^42, 43^. However, the phagocytic ability of zebrafish B cells appears to be negligible, as we show here that they failed to uptake intraperitoneally-injected latex beads, in contrast to professional phagocytes such as macrophages and neutrophils. Notably, while phagocytosis has been proposed as a common feature of teleost B cells^34^, only a few studies so far have explored the phagocytic properties of zebrafish B cells, with contradicting results. Over the course of characterizing B cell ontogeny through the use of new fluorescent transgenic reporter lines, our group initially reported no significant phagocytosis of pHrodo-labeled *E.coli* or latex beads by zebrafish IgM-expressing B cells, either *in vivo* or *in vitro*^26^. In another study by Zhu et al., internalization *in vitro* of soluble KLH by zebrafish B lymphocytes led the authors to consider these cells as phagocytic^44^. However, endocytosis, rather than phagocytosis, is likely to be the main pathway for B cell acquisition of such small antigens^45^. In support of this, the same study found that B cells poorly engulfed large bacteria^44^, whose internalization more likely relies on active phagocytosis^45^. As the phagocytic potential in teleost fish varies between different species and greatly depends on the anatomical source of B cells as well as the nature of the ingested particles^42, 46–49^, it could be hypothesized that zebrafish B cell populations from different compartments have distinct phagocytic capabilities. On the other hand, it is also possible that this feature was lost in zebrafish in the course of evolution, in line with the specific characteristics that separate many bony fish. Further investigation, out of the scope of the present study, will be required to address these important questions. Importantly, the poor phagocytic activity observed in B lymphocytes does not exclude a possible conservation of mpeg1 microbicidal properties within the zebrafish B cell lineage, as B lymphocytes can also be infected by pathogenic bacteria that can enter the cell through phagocytosis-independent mechanisms^50, 51^. It will be interesting to test this hypothesis using live invasive bacterial pathogens.

To conclude, our study unambiguously reveals that *mpeg1.1* does not constitute an ideal marker to track macrophages in the adult zebrafish. While the new observations made here do not complicate studies undertaken at ages before B-lymphocytes appear (at around three weeks of age), it is possible that due to misinterpretation, some of the conclusions drawn in previous reports relying on the specificity of mpeg1.1 expression in adults may need reassessment. For example, a recent study reported the existence of *irf8*-independent macrophages in the gut^52^. Although we cannot exclude that a subpopulation of intestinal macrophages may still develop in the absence of a functional *irf8*, based on our analyses it is more likely that these cells are in fact B-cells. This hypothesis, which remains to be fully explored, is also supported by the severe reduction in macrophage-specific *C1q* genes reported in the gut of *irf8* mutants^52^.

Although the two hematopoietic *mpeg1.1^+^* cell populations can be theoretically discriminated in adult tissues by flow cytometry based on the combination of their light-scatter characteristics and different levels of *mpeg1.1* expression (high for macrophages and low for B lymphocytes), it would be interesting to assess other markers such as *csf1ra* and *mfap4*, for which fluorescent reporters are available^53, 54^. While their specificity in adults remains to be determined, these lines may represent possible alternatives to the use of *mpeg1.1*-driven transgenic lines for the study of macrophages in adult fish. Nevertheless, since none of these tools are ever specific, we would caution investigators working on adult immune cells not to rely exclusively on transgene expression for cell characterization. Rather, we would advise to always complement these analyses with cell isolation and gene expression approaches that will faithfully identify the cells of interest.

## AUTHORSHIP

J.Y.B. and V.W. designed the research and directed the study. G.F., E.G., M.R. and M.M performed zebrafish experiments. S.I. and D.M.L. performed single cell RNA sequencing analysis. J.Y.B. and V.W. wrote the manuscript with comments from all authors.

## ACKNOWLEDGEMENTS

We thank the following for their support and contributions: J-M Vanderwinden and the LiMiF for technical support with confocal imaging, Christine Dubois for help with flow cytometry and Marianne Caron for technical assistance, and members of the Wittamer and Bertrand labs for critical comments and suggestions. V.W. is an investigator of WELBIO and is also supported by grants from the Fonds National de la Recherche Scientifique (FNRS) and The Minerve Foundation. J.Y.B. is funded by the Swiss National Fund (31003_166515). D.M.L. is supported by R24 OD016761 and R01 CA211734 from the National Institutes of Health. G.F. received funding from a FNRS fellowship (Research Fellow).

