## Supplemental Data for "The *macrophage-expressed gene* (*mpeg*) *1* identifies a subpopulation of B cells in the adult zebrafish"

Figure S1

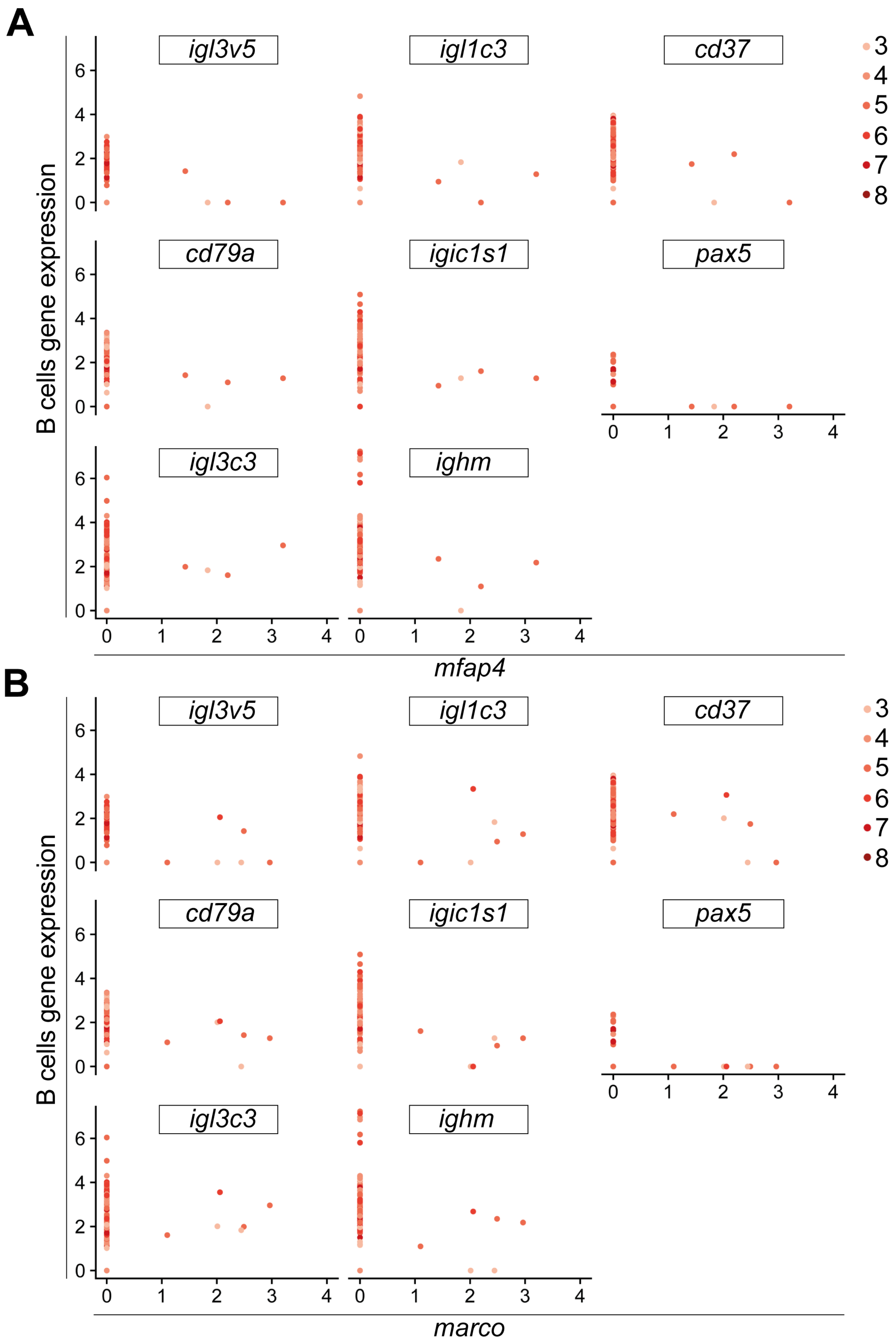
A
igl3v5
igl1c3
cd37
cd79a
igic1s1
pax5
igl3c3
ighm
B cells gene expression
mfap4
3
4
5
6
7
8
B
igl3v5
igl1c3
cd37
cd79a
igic1s1
pax5
igl3c3
ighm
B cells gene expression
marco
3
4
5
6
7
8

**Table 1. qPCR primers used throughout the paper**

| Gene | Forward Primer | Reverse Primer |
|---|---|---|
| *ef1α* | GAGAAGTTCGAGAAGGAAGC | CGTAGTATTTGCTGGTCTCG |
| *mpeg1.1* | CCCACCAAGTGAAAGAGG | GTGTTTGATTGTTTTCAATGG |
| *csf1ra* | ATGACCATACCCAACTTTCC | AGTTTGTTGGTCTGGATGTG |
| *marco* | ACGACAGCTTCGATAATTTG | AAAATACTGCTCTCGGTTCC |
| *cd45* | AGTTCCTGAAATGGAAAAGC | GCACAGAAAAGTCCAGTACG |
| *cldnh1* | TTACAACCCGTTACTGCCCG | TGCCGGCTTGTACTTCTTCT |
